# The role of preadaptation, propagule pressure and competition in the colonization of new habitats

**DOI:** 10.1101/697342

**Authors:** Adriana Alzate, Renske E. Onstein, Rampal S. Etienne, Dries Bonte

## Abstract

To successfully colonize new habitats, organisms not only need to gain access to it, but also need to cope with the selective pressures imposed by the local biotic and abiotic conditions. The number of immigrants, the preadaptation to the local habitat and the presence of competitors are important factors determining the success of colonization. Here, using an experimental set-up, we test the combined effect of propagule pressure, preadaptation and interspecific competition on the colonization success of new habitats using the two-spotted spider mite (*Tetranychus urticae*) as our model system and the red spider mite (*Tetranychus evansi*) as a competitor. Our results show that propagule pressure and preadaptation positively affect colonization success. More successful populations reach larger final population sizes either by having higher per capita growth rate (due to preadaptation effect) or by starting a population with a larger number of individuals. Although populations are more successful colonizing non-competitive environments than competitive ones, propagule pressure and preadaptation counteract the negative effects of competition, promoting colonization success. Our results show the importance of propagule pressure and preadaptation to cope both with the exigencies of the new environment and the community context for successful colonization of new habitats.

## INTRODUCTION

It is well known that new habitats provide novel ecological opportunities, potentially facilitating speciation and diversification (Simpson 1953, Carson & Templeton 1984, Onstein *et al*. 2014, Delaux *et al*. 2015). However, colonization of new habitats very likely relies on the interplay between ecological (e.g. competition, dispersal) and evolutionary (e.g. adaptation) processes that results in complex ecological-evolutionary dynamics. Although some of these processes (dispersal, adaptation and competition) have been studied at large spatial and temporal (evolutionary) scales (Donoghe 2008, Edwards & Donoghe 2013, Onstein et al. 2014), an understanding how these factors interact to affect colonization success of new habitats is still lacking.

For individuals to colonize new habitats, firstly they need to physically access it (via dispersal). Second, the number of individuals arriving in a colonization event (referred to as propagule pressure) affects the probability whether the colonization is successful (Maron 2006, Simberloff 2009). A high propagule pressure increases the chance that some of the immigrants can establish in the new habitat, for instance by having the right genetic makeup. Also, it reduces the chance of extinction, which is more likely to occur in small population sizes due to Allee effects, founder effects, genetic (inbreeding, drift) and demographic stochasticity (Ellstrand & Ellam 1993, Newman & Pilson 1997, Dressler et al. 2019). Third, in order to establish populations and further expand after arrival, individuals need to cope with the ecological context of the new habitat, i.e. the new abiotic conditions (which might differ from the ones in the original environment), and competitors present, resulting in availability of unexploited resources (‘empty niches’) (Simpson 1953). Preadaptation to the new habitat (i.e. genetic or phenotypic traits that have evolved or show phenotypic plasticity to cope in the new environment) may increase the chance of colonization success (Dlugosch & Parker 2007, Kalburge *et al*. 2014, Hamilton *et al*. 2015). Furthermore, these preadaptations may also help individuals to cope with competition from the receiving community. Preadaptation of herbivores to a new host plant may involve the development of new metabolic pathways to metabolize anti-herbivory components and increase resource acquisition and a more efficient exploitation of resources (Dermauw *et al*. 2018, Rane *et al*. 2019, Van Leeuwen & Dermauw, 2016). As a consequence, adaptation to a type of habitat (e.g. a type of plant) would increase fitness in that particular habitat and population growth rate (Hendry 2019, Strauss 2014). In a competitive environment, an increase in population growth rate implies also a change in the competition coefficients, which might bring some advantages to the more adapted individuals and likely increasing their competitive abilities. Therefore, preadaptation might positively affect colonization success by avoiding competitive exclusion. Even though these pre-requisites for colonization success are well known, the relative importance of physical access to the habitat, preadaptation and competition to determine success in new habitats, remains unknown.

Here, we experimentally tested the importance of propagule pressure (number of immigrants in a single dispersal event), preadaptation and competition on colonization success in novel environments. We used the two-spotted spider mite (*Tetranychus urticae*) as our study system. *T. urticae* individuals from different populations were introduced to a novel host plant (tomato), to which some of the populations were pre-adapted and others were not. Competitive environments were created by including competitive mites (*Tetranychus evansi)* in the novel environment. Propagule pressure was examined by varying the initial number of immigrants to the new habitat. Population size and per capita growth rate after one generation in the novel environment was used a proxy of colonization success. We specifically tested three hypotheses: (H1) higher propagule pressures will positively affect population sizes but not per capita growth rate, (H2) preadapted individuals will more rapidly increase in population size and attain higher per capita growth rate in new habitats than less adapted individuals; and (H3) competition will reduce population sizes and population sizes irrespectively of propagule pressure or preadaptation. Our results show that large propagule pressures, preadaptation, and low competition are pre-requisites of successful colonization of novel habitats.

## METHODS

### Study species

The two-spotted spider mite *Tetranychus urticae* Koch, 1836 (Acari, Tetranychidae) is a generalist herbivore that feeds on a wide variety of host plants (Gotoh *et al*. 1993, Bolland *et al*. 1998). Because of its small body size (female size about 0.4mm length), high fecundity (1-12 eggs/day) and short developmental time (11-28 days; Nacimiento de Vasconcelos *et al*. 2008), *T. urticae* is an ideal model organism for mesocosm experiments on adaptation (Gould 1979, Fry 1990, Agrawal 2000, Egas & Sabelis 2001, Magalhaes *et al*. 2007, Kant *et al*. 2008, Bonte *et al*. 2010, Alzate *et al*. 2017, 2019). Moreover, its biology has been widely reported and its genomics is well-known (Grbić *et al*. 2011). All populations used in this study were derived from the London strain, originally collected from the Vineland region in Ontario, Canada (Grbić *et al*. 2011). We used 3 populations of *T. urticae* that differ in their level of adaptation to tomato (the novel environment used in this study): non-adapted, medium adapted and highly adapted. The non-adapted population has been reared on bean plants (*Phaseolus vulgaris* variety “prelude”) for more than 6 years. Both the medium adapted and the highly adapted populations were derived from the non-adapted population, but medium adapted populations have been obtained by rearing four different populations on tomato plants for about 20 generations (Alzate *et al*. 2017), and the highly adapted population has been reared on tomato plants for more than 100 generations. These populations differed in their fitness, measured as fecundity (number of eggs as a proxy of fitness), on tomato plants, suggesting differences in their adaptation to the tomato host plant (Alzate *et al*. 2017, Fig. 1).

**Fig. 1.**
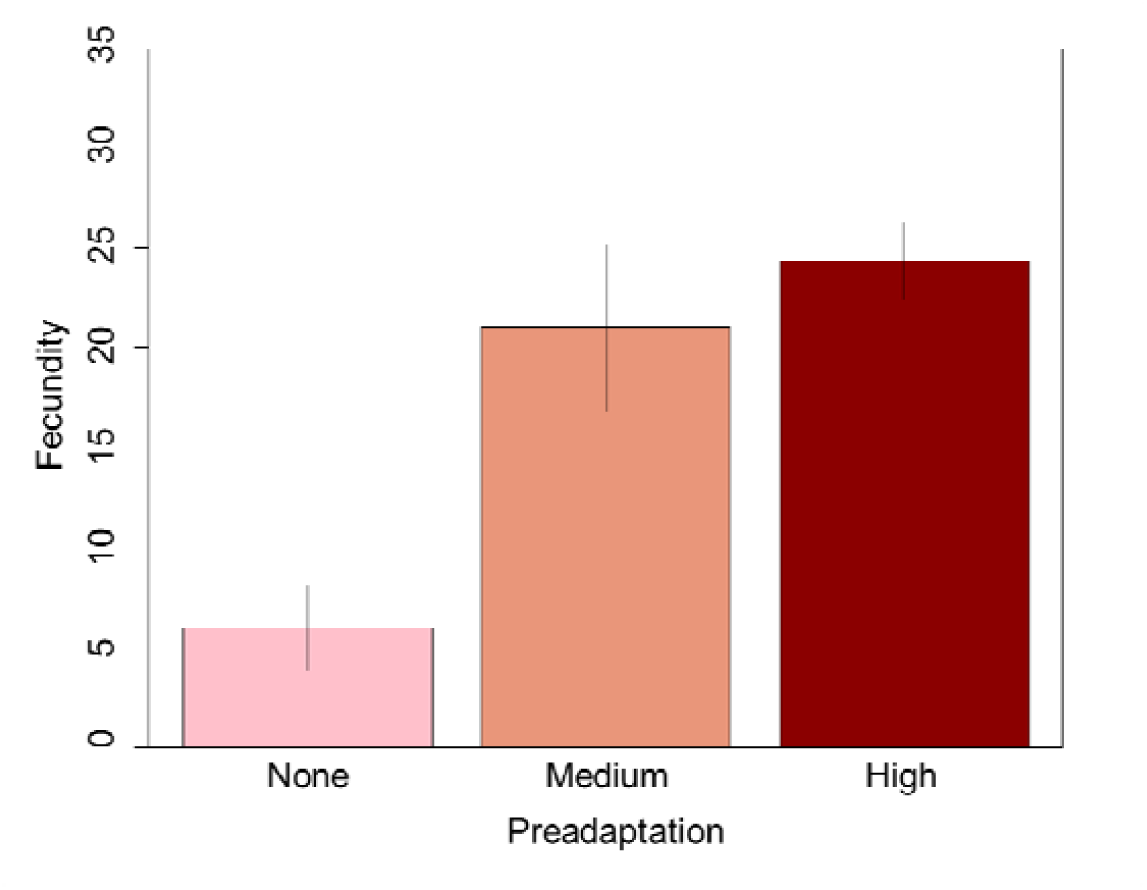
From Alzate *et al*. 2017: population used in this study show different levels of adaptation to tomato plants. Fecundity (number of eggs a single female laid during 6 days) was considered as a proxy of adaptation to tomato plants. The population with no preadaptation has never been exposed to tomato (only reared on bean plants), populations of females with medium preadaptation have been exposed to tomato plants for 20 generations (1 generation ∼ 13 days), and the population with high preadaptation has been exposed to tomato plants for more than 100 generations.

As a competitor, we used the red spider mite *Tetranychus evansi* Baker and Pritchard, 1960 (Acari, Tetranychidae), which is a specialist herbivore of (mainly) Solanaceae (incl. tomato). Adult females are easily distinguishable from *T. urticae* as they show a characteristic red coloration and are slightly larger (0.5 to 0.6mm length). Fecundity ranges from 10 to 14 eggs per day (Navajas *et al*. 2013) and development time can vary from 6.3 to 13.5 days, depending on the environmental temperature and host (Bonato 1999).

### Experiments

We performed two experiments: in one we tested the effects of propagule pressure and competition on colonization success and abundance and in the other we tested for the effects of preadaptation and competition on colonization success and abundance. Before each experiment we removed epigenetic effects (juvenile and maternal effects) by collecting individual females from each population (non-adapted, medium adapted and highly adapted) and rearing them separately in a common garden for 2 generations (Kawecki *et al*. 2012, Magalhães *et al*. 2011). The common garden consisted of a 5cm diameter bean leaf disk (per female) on cotton wool soaked in distilled water. All individuals derived from a single female are therefore considered an iso-female line and each line was used as a replicate for the experiments performed in this study.

#### Propagule pressure and competition

To test the effect of propagule pressure and competition on colonization success, we used the highly adapted population. We tested three levels of propagule pressure (3, 5, 10 individual adult female mites), and the presence or absence of competition with *T. evansi* (3 individuals). Per iso-female line, we placed adult female mites (3, 5 or 10) on either a complete (four weeks old) tomato plant with or without competition. In total we tested six treatment combinations with eight replicates (8 iso-female lines) for treatments with propagule pressure of 3 individuals and five replicates (5 iso-female lines) for treatments with propagule pressure of 5 and 10 individuals (Fig. S1 in Supplementary material).

#### Preadaptation and competition

To evaluate the effect of preadaptation and competition on colonization success to new environments, we placed 3 adult females from each adaptation treatment (and iso-female line) either on a complete (four weeks old) tomato plant without cohabitants (no competition treatment) or on a complete tomato plant together with 3 females of *T. evansi* (competition treatment). We tested 3 preadaptation and 2 competition levels, for a total of 6 treatment combinations. We used 8 replicates (iso-female lines) for treatments with non-adapted and highly adapted populations and 12 replicates for the treatment with medium adapted populations. Medium adapted populations have more replicates because we collected females from four independent populations, whereas for the non-adapted and highly adapted treatments, females came from a single population (Fig. S2 in Supplementary material).

For both experiments, plants were maintained in a climate regulated room at 25 ± 0.5°C with a 16/8h light/dark regime for 15 days. Population sizes and growth rate of *T. urticae* were recorded after 15 days (one generation). To estimate population sizes, we counted all adult female mites present on each complete tomato plant. Juveniles and males were not included in the counting because their small size made their detection with the naked eye difficult. Real population sizes are therefore larger than the ones presented in this study, when accounting for juveniles and males (S2 in De Roissart *et al*. 2015). Per capita growth rate was calculated by first subtracting the initial number of females (e.g. propagule pressure) from the final number of females after one generation, then dividing this by the initial number of females.

### Statistical analyses

#### Propagule pressure and competition

to test the effect of propagule pressure and competition on per capita growth rate and population size after one generation, we used a general linear mixed model. Propagule pressure (with three levels: 3, 5 and 10 female mites) and competition (with two levels: competition with T. evansi and no-competition) were considered as fixed categorical factors. Because females coming from the same iso-female line (Fig. S1 in Supplementary material) might respond similarly to a treatment than females that are not related, iso-female line was considered as random factor. Per capita growth rate and final population size (number of adult females after one generation) were considered as response variables. To correct for the initial differences in population sizes on final population sizes, we subtracted the initial number of immigrants (3, 5 or 10 female mites) from the final population size. Model selection (for both the random and fixed part) was performed using a stepwise removal of non-significant effects based on log-likelihood ration test until only significant effects remained. Post-hoc tests were performed to test for differences between the least square means of treatments using the *difflsmeans* function from the package lmerTest (Kuznetsova et al. 2016). Degrees of freedom were calculated with a Satterthwaite’s approximation.

#### Preadaptation and competition

To test the effect of preadaptation and competition on per capita growth rate and final population size, we used a general linear mixed model where preadaptation (3 levels: no adapted, medium adapted and highly adapted) and competition (2 levels: competition and no competition) were considered as fixed factors. Iso-female line was considered as random factor. Model selection (for both the random and fixed part) was performed using a stepwise removal of non-significant effects based on a log-likelihood ratio test until only significant effects remained. Per capita growth rate and final population sizes were log-transformed to meet the assumption of normal distribution of model residuals. Post-hoc tests were performed to test for differences between the least square means of treatments using the *difflsmeans* function from the package lmerTest (Kuznetsova et al. 2016). Degrees of freedom were calculated with a Satterthwaite’s approximation.

All analyses were performed in R version 3.5.3 (R core Team 2019) and the R packages lme4 (Bates *et al*. 2015), MuMIn (Barton 2018), lsmeans (Lenth 2016), lmerTest (Kuznetsova *et al*. 2017), multcomp (Hothorn *et al*. 2008).

## RESULTS

### Propagule pressure and competition

Final population size was best explained by the additive effects of propagule pressure and competition (Table 1, see Table S1 in Supplementary material for model selection). While propagule pressure positively affected population size of *T. urticae*, both with and without competition, competition always exerted a negative effect on population size irrespective of propagule pressure (Fig. 2a). Populations receiving a higher number of propagules (10 females) attained larger population sizes after one generation than populations receiving fewer propagules (3 females).

**Table 1.**
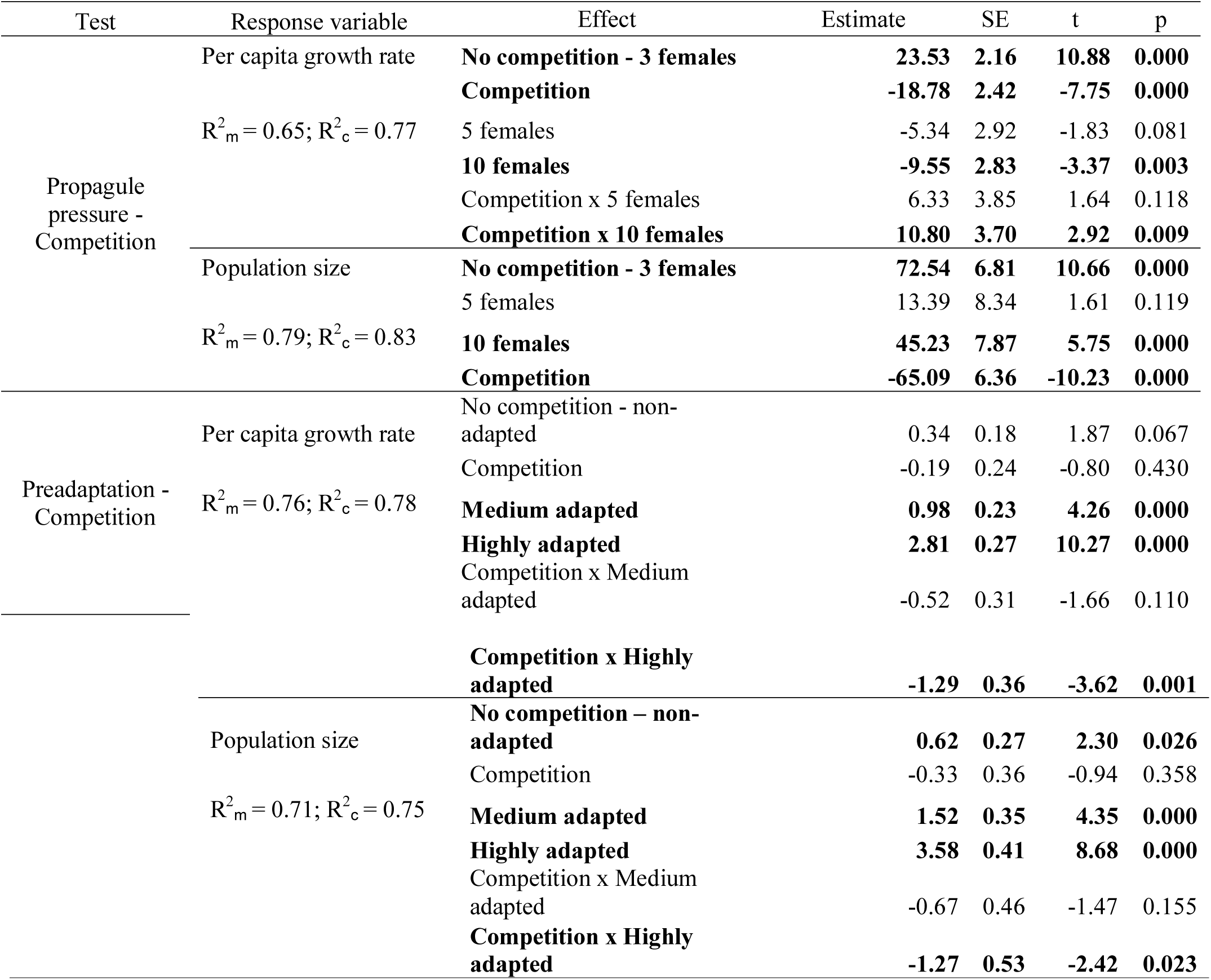
Summary of final statistical models testing 1) the effects of propagule pressure and competition and 2) the effects of preadaptation and competition on per capita growth rate and population size of *T. urticae* when colonizing a novel habitat. Marginal (fixed effects only) and conditional (fixed and random effects) coefficients of determination for the final models are shown (R^2^_*m*_, R^2^_*c*_ respectively). Estimates for models testing preadaptation and competition are log-transformed.

**Figure 2.**
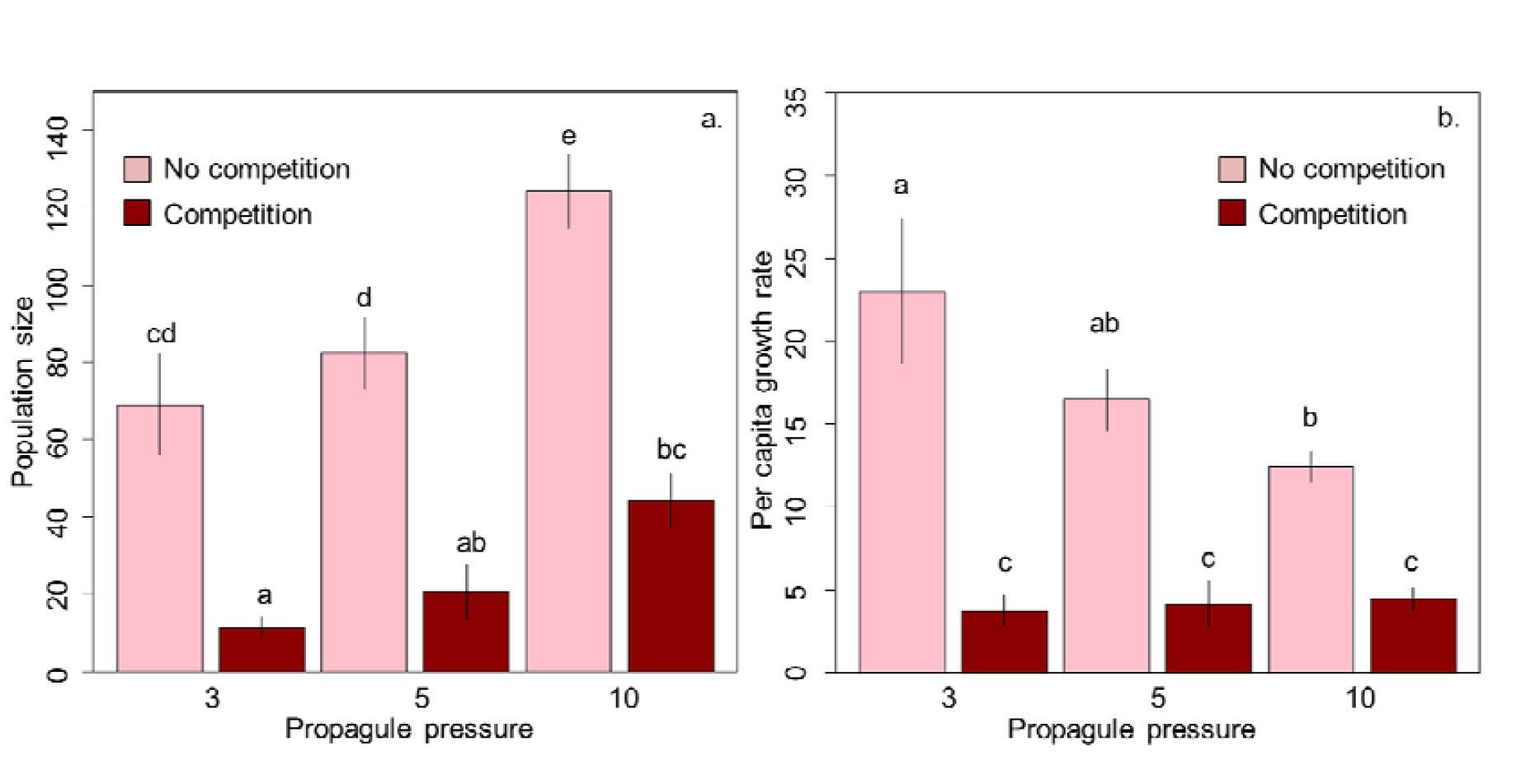
Effect of propagule pressure (number of *T. urticae* colonizers) on population size (a) and per capita growth rate (b) of *T. urticae*. Graph shows the mean population size and standard error for the raw data. Letters show significant differences between treatments.

Per capita growth rate was best explained by both the additive and interaction effects of propagule pressure and competition (Table 1, Table S1 in Supplementary material for model selection). While competition always affected growth rate negatively, the effect of propagule pressure depended on the competitive environment (Table 1). In a no competitive environment, propagule pressure exerted a negative effect on per capita growth rate, whereas in a competitive environment propagule pressure did not have an effect (Fig. 2b).

### Preadaptation and competition

Population size was best explained by the additive and interaction effects of preadaptation and competition (Table 1, see Table S1 in Supplementary material for model selection). Populations that co-occurred with *T. evansi* were significantly smaller than populations without the competitor for populations with medium and high preadaptation (Fig. 3b). Populations with no preadaptation had the lowest population size in both competitive and non-competitive environments, and populations with high preadaptation in non-competitive environments had the largest population sizes (Fig. 3b).

**Fig. 3.**
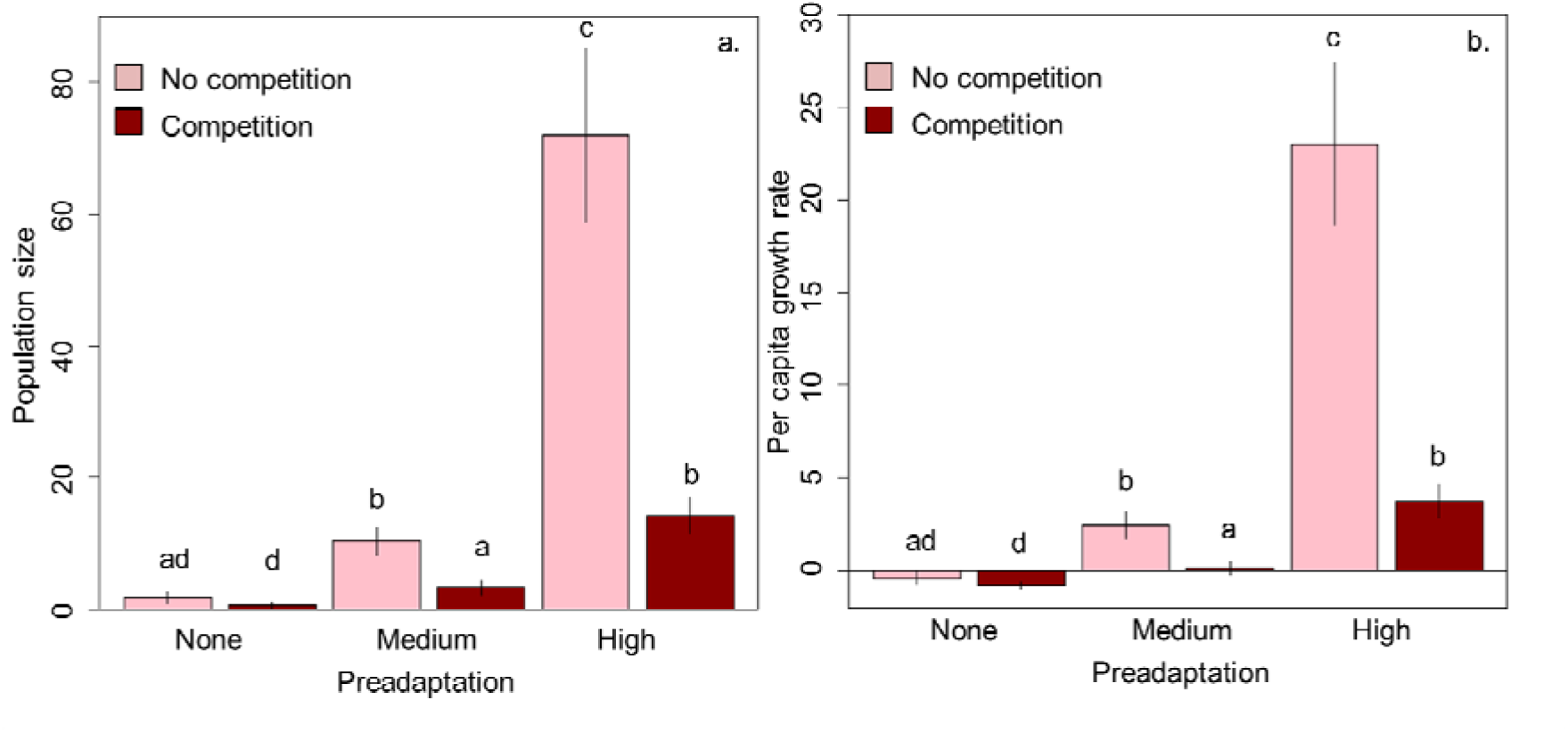
Effect of preadaptation and competition on population size (a) and per capita growth rate (b) of *T. urticae* after one generation on a novel environment. Letters show the significant differences between treatments.

Per capita growth rate was best explained by the additive and interaction effects of preadaptation and competition. Preadaptation affected growth rate positively, whereas competition affected growth rate negatively (Fig. 3a). Populations without any preadaptation showed the lowest growth rate, both in competitive and non-competitive environments, whereas populations with high preadaptation in non-competitive environments showed the highest growth rate (Fig. 3a).

## DISCUSSION

Using microcosms experiments, we investigated the effects of propagule pressure, preadaptation and interspecific competition on colonization success of *T. urticae* to novel environments. Our results show that preadapted populations that arrive in low competitive environments with high propagule pressure are more likely to successfully colonize new habitats. Nevertheless, preadaptation and propagule pressure benefit successful colonization differently. While preadaptation positively affects population sizes by increasing per capita growth rate, propagule pressure increases population sizes due to the additive effect of starting with a large population size. In either way both propagule pressure and preadaptation help the colonizing population cope with competition in the new environment.

Propagule pressure has been widely studied using empirical, observational and experimental approaches mainly because of its relevance on invasion biology and its role on colonization success of invaders (Blackburn & Duncan 2001, Simberloff 2009, Lockwood *et al*. 2005, Cassey *et al*. 2018). In general, propagule pressure has been shown to be positively related to colonization and establishment success (Maron 2006, reviewed in Cassey *et al*. 2018), and our results are in line with these findings. Our experiment is the first, to our knowledge, that assessed the combined effects of propagule pressure (abundance) and interspecific competition on colonization success. Our results show that the ecological context in which individuals arrived (competition vs. no interspecific competition) strongly influenced population size. While we show that propagule pressure positively affected population size, competition counteracted this effect (Fig. 2b). In a competitor-free environment, propagule pressure negatively affected per capita growth rate, whereas it had no effect in a competitive environment. This strongly suggests that intraspecific competition increases with propagule pressure, which in turn hinders growth. In a competitive environment, interspecific competition reduces population sizes to a degree that intraspecific competition may be of less importance. Our results suggest that, in the presence of interspecific competition, the negative effect of propagule pressure on growth rate is absent. Nevertheless, propagule pressure does have a positive effect on the total population size. Since larger populations are less affected by demographic stochasticity and therefore less likely to go extinct than smaller populations (Alzate *et al*. 2019), total population size may be a more relevant indicator for colonization success than growth rate.

Besides the ecological context, the nature of the new habitat can determine the fate of the colonizing individuals. The outcome would depend on the relative differences between the old and the new habitat, and the degree of niche-conservatism of arriving individuals. In other words, immigrants that come from a very different habitat might be less able to cope with the new habitat than individuals that come from a similar habitat which are likely preadapted. Studies on invasion ecology suggest that preadaptation (or proxies such as phylogenetic relatedness to the local species) can increase the chance of successful colonization (Li *et al*. 2015). However, Maron (2006) shows that latitude (their surrogate for preadaptation) does not affect colonization success. Nevertheless, to truly assess the importance of preadaptation, it is important to evaluate the environmental conditions that may require adaptations and thus constrain survival and fitness of organisms based on their morphology, physiology and reproduction (Colautti & Barret 2013). In our study, mites were preadapted to a new host plant (tomato) on which survival and fecundity is much lower in absence of preadaptation (Fig. 1, Alzate *et al*. 2017), probably due to herbivore-induced plant defenses that hampers feeding and reproduction (Kant *et al*. 2015, Godihno *et al*. 2016). Here, we show that preadaptation to the new habitat is key for successful colonization, as it increases population growth rates and total population sizes. Although preadapted populations are, in absolute terms, more affected by competition, they have a higher growth rate independently of the competitive environment. Therefore, preadapted populations attain larger population sizes, which reduces extinction. Preadaptation helps individuals to cope with competitive by increasing their competitive abilities (e.g. more efficient resource consumption). For example, in *Drosophila* species, it has been shown that adaptation to the abiotic environment is the most important component to increase competitive ability (Joshi & Thompson 1996). In our experiment, the most adapted populations not only have larger growth rates and larger final population sizes (Fig. 3), they also exerted a higher competitive effect on the competitor species, *T. evansi* (Fig. S3a in Supplementary material) and displayed a higher overall competitive ability (Fig. S3b in Supplementary material).

## CONCLUSION

Understanding the factors that affect species colonization success in novel habitats is of great importance given the biodiversity crisis we are currently facing. Habitat fragmentation and transformation have forced populations into novel habitats. Inability to successful colonize those habitats may lead species to extinction as populations become more isolated attaining smaller population sizes (Fahrig 1997, Wiegand *et al*. 2005). In this study, we experimentally tested how propagule pressure, preadaptation and competition affect colonization success in novel habitats. Our results confirm the intuitively evident hypothesis that propagule pressure and preadaptation positively affect colonization success. In competitive environments, however, colonization success is reduced, and to successfully colonize habitats that lack ‘empty niches’ (which is likely the most common scenario), it is important to be either preadapted or to start a population with a large number of colonizing individuals (higher propagule pressure).

## ACKNOWLEDGMENTS

We thank Karen Bisschop, Pieter Vantieghem and Angelica Alcantara for their help during the experiments. RSE thanks the Netherlands Organisation for Scientific Research (NWO) for financial support through a VICI grant. AA was funded by the Ebbo Emmius Fund from the University of Groningen, AA and DB were funded by BelSpo IAP project ‘SPatial and environmental determinants of Eco-Evolutionary DYnamics: anthropogenic environments as a model’; DB and RSE by the FWO research community ‘An eco-evolutionary network of biotic interactions’. AA and REO were financially supported by the German Center for Integrative Biodiversity Research (iDiv) Halle-Jena-Leipzig. Drawing of tomato plant, mites and beans were obtained from the noun project (https://thenounproject.com) and the artists Michael Zick Doherty, Vectors Market and Xela Ub.

