## Supplementary material for "The role of preadaptation, propagule pressure and competition in the colonization of new habitats"

**Fig. S1.** Design of propagule pressure - competition experiment. We removed epigenetic effects (juvenile and maternal effects) by collecting individual females from the highly adapted population (reared on tomato plants). We place each female separately in a common garden for 2 generations. The common garden consisted of a 5cm diameter bean leaf disk (per female) on cotton wool soaked in distilled water. All individuals derived from a single female are therefore considered an iso-female line and each line was used as a replicate for the experiments performed in this study. Per iso-female line, we placed adult female mites (propagule pressure: 3, 5 or 10 females) on a complete (four weeks old) tomato plant either with or without competition. In total, we tested six treatment combinations with eight replicates (8 iso-female lines) for treatments with propagule pressure of 3 individuals and five replicates (5 iso-female lines) for treatments with propagule pressure of 5 and 10 individuals.

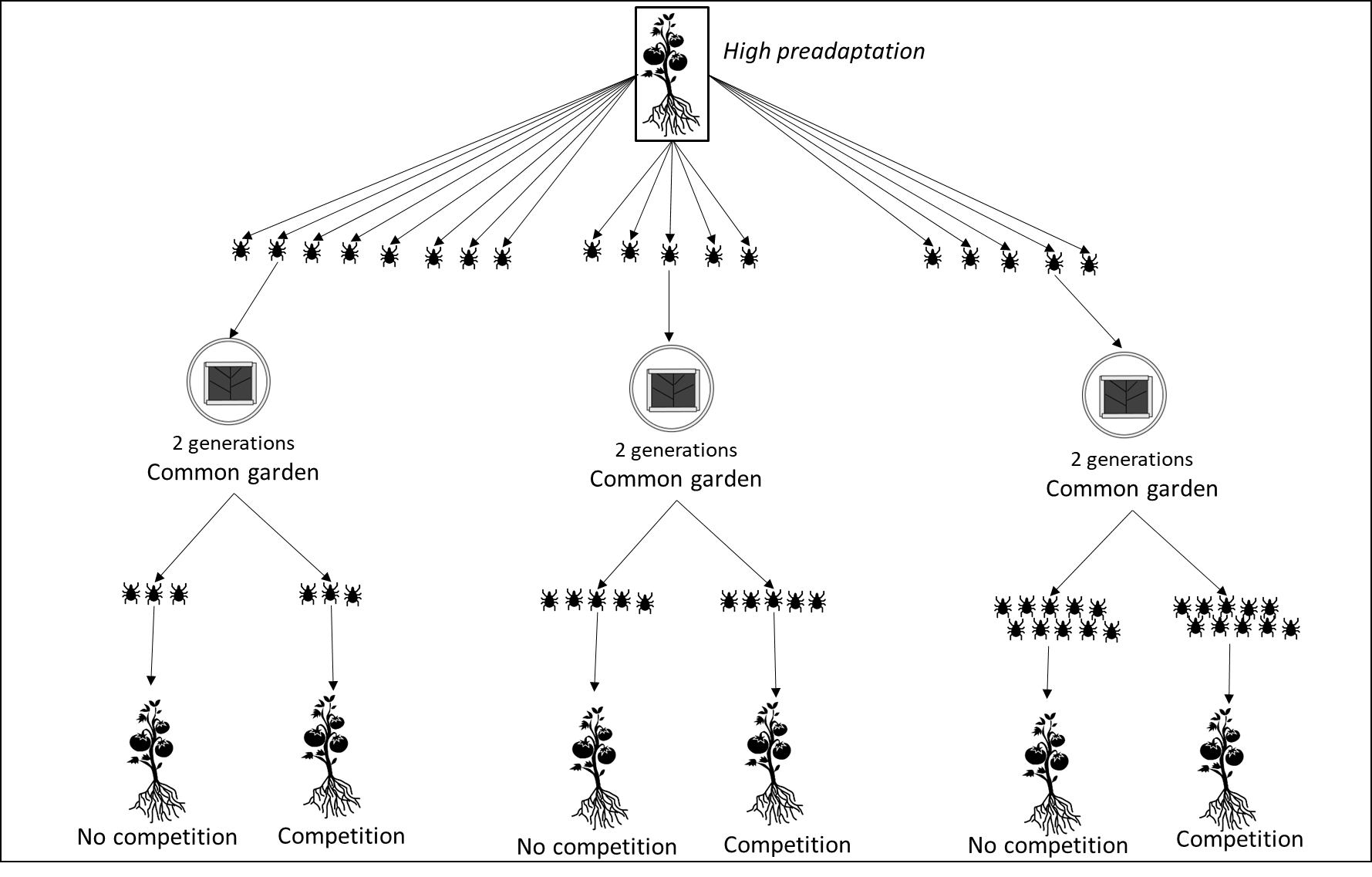

**Fig. S2.** Design pre-adaptation - competition experiment. We removed epigenetic effects (juvenile and maternal effects) by collecting individual females from each population (non-adapted, medium adapted and highly adapted). We place each female separately in a common garden for 2 generations. The common garden consisted of a 5cm diameter bean leaf disk (per female) on cotton wool soaked in distilled water. We placed 3 adult females from each adaptation treatment (and iso-female line) on a complete (four weeks old) tomato plant either without competition or on a complete tomato plant together with 3 females of *T. evansi* (competition treatment). We tested 3 preadaptation and 2 competition levels, for a total of 6 treatment combinations. We used 8 replicates (iso-female lines) for treatments with non-adapted and highly adapted populations and 12 replicates for the treatment with medium adapted populations. Medium adapted populations have more replicates because we collected females from four independent populations, whereas for the non-adapted and highly adapted treatments, females came from a single population.

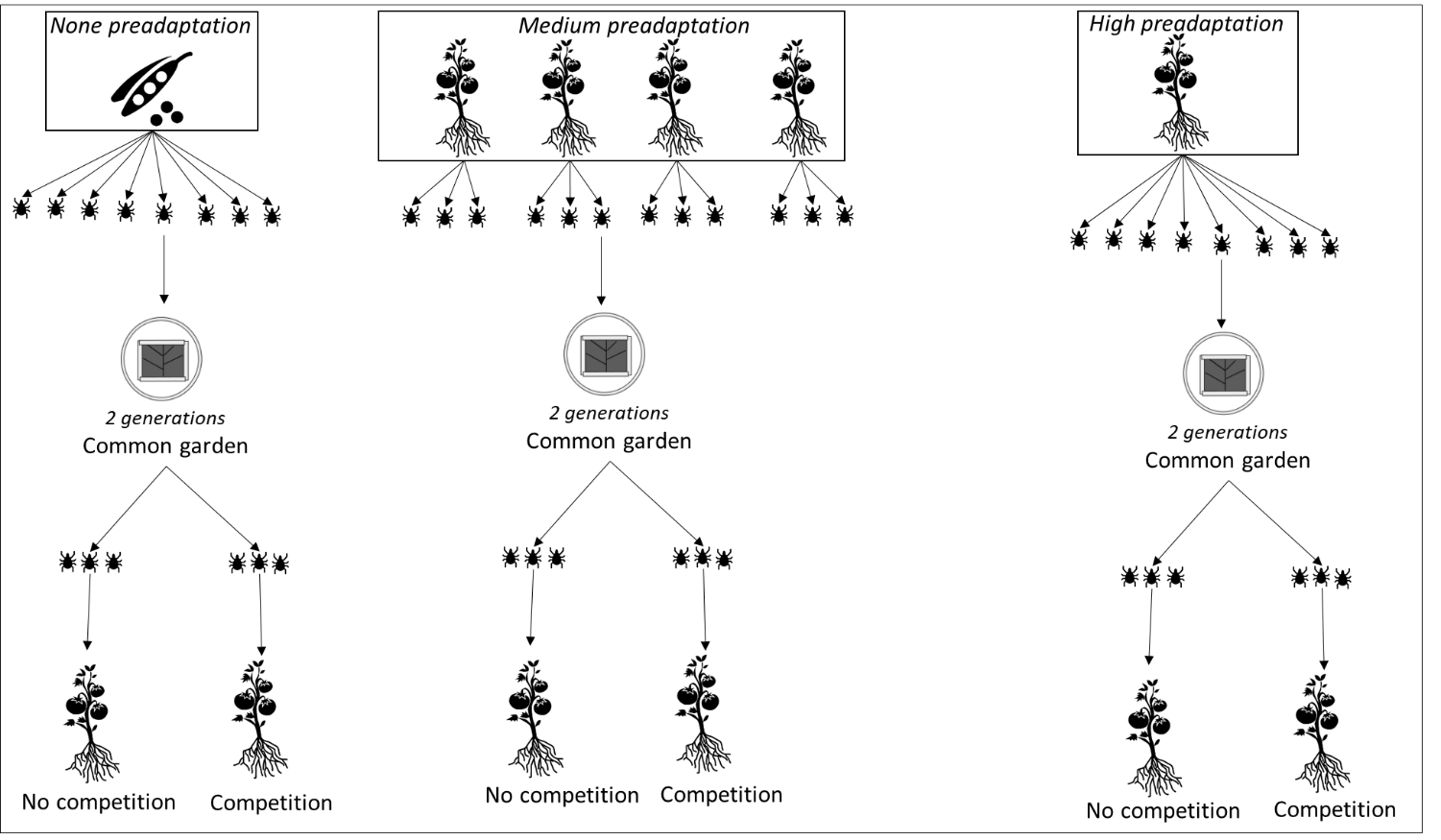

**Table S1.** Model selection for the effect of propagule pressure/pre-adaptation and interspecific competition on per capita growth rate/ population size. The selection was performed a backward step‐wise removal of non‐significant effects (> 0.05), starting with the interaction terms

| Test | Model comparison | *df* | AIC | BIC | log Lik | deviance | Chi ^2^ | Chi df | *p* |
| --- | --- | --- | --- | --- | --- | --- | --- | --- | --- |
| per capita growth rate ~ Propagule pressure & Competition | per capita growth rate ~ Competition + Propagule pressure + (1 \| iso-female line) | 6 | 214.97 | 223.95 | -101.49 | 202.97 |  |  |  |
|  | **per capita growth rate ~ Competition * Propagule pressure + (1 \| iso-female line)** | **8** | **209.95** | **221.92** | **-96.97** | **193.95** | **9.02** | **2.00** | **0.011** |
| Population size ~ Propagule pressure & Competition | **Population size ~ Competition + Propagule pressure + (1 \| iso-female line)** | **6** | **297.57** | **306.55** | **-142.78** | **285.57** |  |  |  |
|  | Population size ~ Competition * Propagule pressure + (1 \| iso-female line) | 8 | 298.50 | 310.47 | -141.25 | 282.50 | 3.07 | 2.00 | 0.215 |
| per capita growth rate ~ Preadaptation & Competition | log(per capita growth rate + 2) ~ Competition + Adaptation + (1 \| iso-female line) | 6 | 97.74 | 109.68 | -42.87 | 85.74 |  |  |  |
|  | **log(per capita growth rate + 2) ~ Competition * Adaptation + (1 \| iso-female line)** | **8** | **89.22** | **105.13** | **-36.61** | **73.22** | **12.53** | **2.00** | **0.002** |
| Population size ~ Preadaptation & Competition | log(Population size + 1) ~ Competition + Adaptation + (1 \| iso-female line) | 6 | 135.58 | 147.52 | -61.79 | 123.58 |  |  |  |
|  | **log(Population size + 1) ~ Competition * Adaptation + (1 \| iso-female line)** | **8** | **133.53** | **149.44** | **-58.77** | **117.53** | **6.05** | **2.00** | **0.048** |

**Fig. S3.** a) effect of *T.urticae* preadaptation on *T.evansi* population size. b) effect of preadaptation on the overall competitive ability of *T. urticae*. Overall competitive ability was calculated as the fraction of individuals of *T. urticae* among the total number of individuals (*T.urticae* + *T.evansi*).

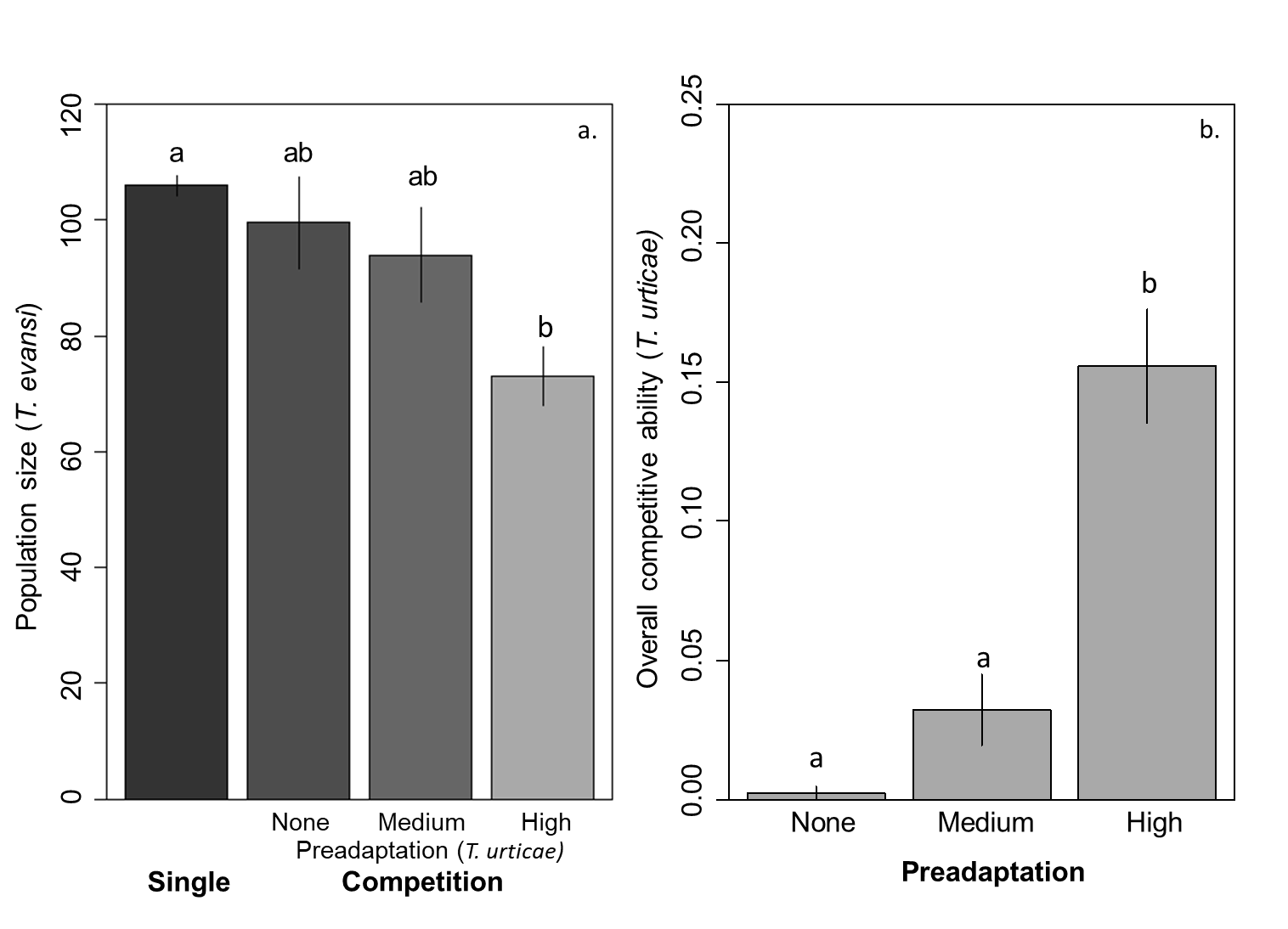
